# The mismatch repair factor Mlh1-Pms1 uses ATP to compact and remodel DNA

**DOI:** 10.1101/2025.01.16.633381

**Authors:** Bryce W. Collingwood, Amruta N. Bhalkar, Carol M. Manhart

## Abstract

In eukaryotes, mismatch repair begins with <u>M</u>ut<u>S h</u>omolog (MSH) complexes, which scan newly replicated DNA for mismatches. Upon mismatch detection, MSH complexes recruit the PCNA- stimulated endonuclease Mlh1-Pms1/PMS2 (yeast/human), which nicks the DNA to allow downstream proteins to remove the mismatch. Past work has shown that although Mlh1-Pms1 is an ATPase and this activity is important *in vivo*, ATP is not required to nick DNA. Our data, using yeast as a model, suggests that Mlh1-Pms1 forms oligomeric complexes that drive DNA conformational rearrangements using the protein’s ATPase activity. Experiments with non-B-form DNA structures, common in microsatellite regions, show that these structures inhibit Mlh1-Pms1’s activities, likely through impeding Mlh1-Pms1-dependent DNA conformational changes. This could explain an additional mode for instability in these regions of the genome. These findings highlight the importance of DNA compaction and topological rearrangements in Mlh1-Pms1’s function and provide insight into how mismatch repair relies on DNA structure to coordinate events.

**GRAPHICAL ABSTRACT:** 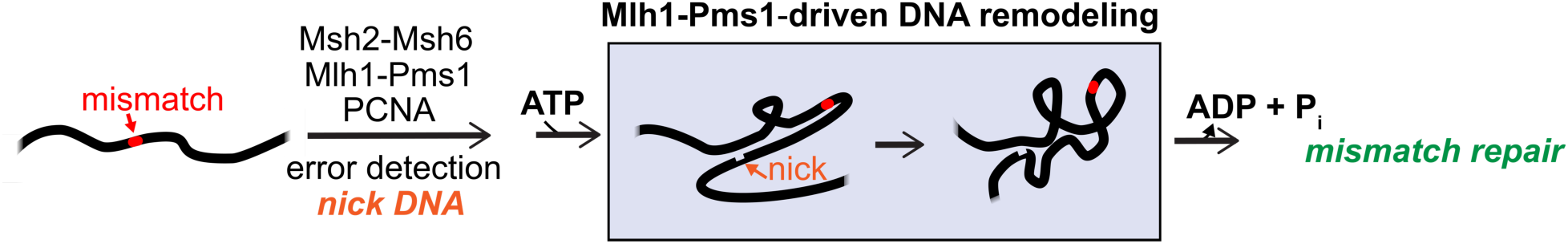

## INTRODUCTION

DNA mismatch repair is a highly conserved biological process crucial for maintaining genomic stability. It primarily corrects mispaired nucleotides that arise from errors during DNA replication or repair, thereby preventing the accumulation of mutations. Defects in this process are associated with Lynch Syndrome, a hereditary cancer syndrome characterized by increased cancer risk due to unrepaired DNA mutations (1). In eukaryotic mismatch repair, mismatches are recognized by either the Msh2-Msh6 or Msh2-Msh3 complex, which have overlapping specificities (2). These MutS homolog complexes recruit the MutL homolog endonuclease (Mlh1-Pms1 in yeast; MLH1-PMS2 in humans and mice). The Mlh1-Pms1/PMS2 complex is stimulated by the replication processivity clamp PCNA to nick one strand of the DNA duplex (3). This nick serves as an entry point for other factors that remove the mismatch through polymerase strand displacement and flap cleavage, or through excision and resynthesis (3, 4).

The Mlh1-Pms1/PMS2 complex is critical for initiating mismatch removal in eukaryotic systems. This protein consists of two homologous subunits, each containing a globular amino-terminal domain important for DNA binding and ATPase activity (5–8). These domains are connected by intrinsically disordered regions to globular carboxy-terminal domains, which contain the primary dimerization site between the Mlh1 and Pms1/PMS2 subunits. The endonuclease active site is primarily located in the carboxy-terminal domain of the Pms1/PMS2 subunit (9).

The Mlh1-Pms1/PMS2 complex is highly dynamic and undergoes significant conformational changes mediated by its intrinsically disordered regions (10–16). Studies on the *E. coli* MutL homodimer, which is homologous to Mlh1-Pms1/PMS2, suggest that ATP binding induces conformational changes that allow the amino-terminal domains to interact like a clamp, with ATP mediating the opening and closing of the protein’s structure (10, 11). Similar dynamics have been observed in yeast and human systems, where asymmetric ATPase activities in the Mlh1 and Pms1/PMS2 subunits drive sequential and allosteric conformational changes (12, 17–20). Visual evidence from atomic force microscopy studies in yeast and human systems shows that ATP binding leads to large-scale structure changes, with the intrinsically disordered regions condensing to bring the amino-and carboxy-terminal domains closer together (12). However, these conformational changes have primarily been studied in the absence of DNA, leaving their role for DNA-bound states unclear.

ATPase activity not only controls the conformational dynamics of Mlh1-Pms1/PMS2 but is also critical for its *in vivo* function. In yeast, mutations that disrupt ATP binding in either the Mlh1 or Pms1 subunits result in mutation rates similar to those observed in *mlh1Δ* or *pms1Δ* strains, demonstrating the importance of ATPase activity in mismatch repair (17, 18). Interestingly, neither yeast Mlh1-Pms1 nor nucleolytic MutL from *B. subtilis* require ATP to nick DNA *in vitro*, suggesting that ATPase activity might instead modulate dynamics needed for recycling the protein after nicking DNA or recruiting other factors (13, 21, 22). Biochemical studies also indicate that ATP-driven conformational changes in the intrinsically disordered regions occur after endonuclease activation, further supporting the idea that ATPase activity is required for an Mlh1-Pms1 activity distinct from DNA cleavage (13, 14, 16). Despite these insights, the precise mechanisms by which ATPase activity regulates Mlh1-Pms1/PMS2’s functions in mismatch repair and its interactions with DNA remain to be elucidated.

In addition to its dynamic nature, Mlh1-Pms1/PMS2 binds DNA cooperatively, with multiple copies assembling on DNA during mismatch repair (23–29). A role for an Mlh1-Pms1/PMS2 oligomer in DNA mismatch repair has not been clearly established, but may provide an explanation for observations that Mlh1-Pms1/PMS2 nicks have been observed in some systems at distances on the order of a few hundred nucleotides from the mismatch (30, 31). If Mlh1-Pms1/PMS2 nicks distant to the mismatch, multiple copies of Mlh1-Pms1/PMS2 may serve as a communication channel between Msh2-Msh6 bound to the mismatch and the nick site. This is consistent with recent work suggesting that MutL homolog complexes can restrain Msh2-Msh6 at the mismatch, although other models suggest that MutS and MutL may move together (26, 32–35).

Beyond acting as a physical bridge, Mlh1-Pms1/PMS2 oligomers have been proposed to promote DNA-DNA interactions. Although previously suggested for the meiotic homolog Mlh1-Mlh3 using *S. cerevisiae* as a model organism (27), yeast Mlh1-Pms1 complexes have been shown to simultaneously interact with two distinct DNA substrates and that this tethering feature can stimulate Mlh1-Pms1 endonuclease activity (28). These data are consistent with atomic force microscopy work visually suggesting that tracts of Mlh1-Pms1/PMS2 can form on DNA and alter the shape of the substrate (23, 26). How Mlh1-Pms1/PMS2 changes the shape of the DNA in mismatch repair and whether large-scale conformational changes in Mlh1-Pms1/PMS2 mediated by ATP control Mlh1-Pms1/PMS2 oligomers and DNA-DNA associations and conformation remains to be determined.

In this study, using purified Mlh1-Pms1 from *S. cerevisiae*, we observed that Mlh1-Pms1 promotes DNA conformational changes or remodeling *in vitro*, an activity regulated by independent protein-DNA interactions of both subunits and modulated by ATP. We also found that sequence features affecting DNA conformation interfere with Mlh1-Pms1’s ability to promote DNA remodeling, potentially explaining instability in genomic regions containing these sequences. These findings highlight a critical step in mismatch repair and provide insights into how sequence context contributes to genome instability and disease.

## MATERIALS AND METHODS

### Reagents Used in this Study

Wild-type Mlh1-FLAG-Pms1 was used for all experiments unless specifically noted. Mlh1-FLAG-Pms1 was expressed from a strain transformed with pMH8, which was a gift from Thomas Kunkel’s lab and pEAE269, which was a gift from Eric Alani’s lab (16, 36). pEAE269 is pMH1, originally constructed by Thomas Kunkel’s lab, with a FLAG tag inserted at amino acid 499 of Mlh1. The Mlh1-Pms1 ATPase mutants (mlh1N35A-pms1N34A, mlh1N35A-Pms1, and Mlh1-pms1N34A) were generated by Q5 site-directed mutagenesis (NEB) using pMH1 and pMH8 as described previously (22). Yeast RFC and PCNA were expressed and purified as previously reported (37, 38).

Unless otherwise indicated, supercoiled plasmid substrates were pUC18 or pUC19 and relaxed plasmid substrates were topoisomerase relaxed pUC18 or pUC19. pUC18 and pUC19 are identical in size and sequence and only differ by the direction of the multiple cloning site. The relaxed forms were generated by incubating 3.36 pmol of supercoiled pUC18/pUC19 with 30 units of DNA Topoisomerase I (NEB) for 1 hour at 37°C in a 60 μL reaction containing 50 mM potassium acetate, 20 mM Tris-acetate, 10 mM magnesium acetate, 100 μg/mL recombinant albumin, pH 7.9). The topoisomerase was then inactivated by incubating at 80°C for 20 min.

Plasmid substrates containing a non-B-form, nucleotide repeat segment were generated by digesting 3.36 pmol of pUC18 with EcoRI (NEB) and BamHI (NEB). The linear fragment was gel extracted (QIAGEN) and 1.4 nM of the extracted product was then incubated with 2 μM of annealed complementary oligonucleotides (Table S1). 400 units of T4 DNA ligase (NEB) was included and incubated for 30 minutes at 16°C in a final volume of 20 μL. The DNA was then transformed and propagated using DH5*α*cells, miniprepped, and insertion of the desired sequence was confirmed by Sanger sequencing (Azenta).

The formation of non-B-form structures was verified using T7 endonuclease I, a structure-selective endonuclease that acts on non-B-form DNA. To do this, 3.8 nM of miniprepped, supercoiled plasmid was incubated with 100 units of T7 endonuclease I (NEB) in a reaction containing 10 mM Tris-HCl, 50 mM NaCl, 10 mM MgCl_2_, 1 mM DTT, at pH 7.9 for 45 minutes at 37°C. Reaction products were analyzed using a 1% agarose gel stained with 1.5 μg/mL ethidium bromide. To characterize the linearized forms of the substrates, an identical procedure was performed immediately after linearization with BsaI-Hfv2 (NEB), which was performed at 37°C for 60 minutes, followed by heat inactivation of the enzyme according to the manufacturer’s instructions.

### High-Throughput DNA Binding Assays for Measuring Affinity for Different DNA Topologies

A pool of topologically distinct, DNA isomers was generated by incubating 3.36 pmol of pUC19 plasmid with 5 units of DNA Topoisomerase I (NEB) for 1 minute at ambient room temperature in the supplied 1x CutSmart Buffer (NEB), followed by heat inactivation of the enzyme at 80°C for 20 minutes. 20 μL reactions were assembled in Buffer A, containing 20 mM HEPES-KOH pH 7.5, 6% glycerol, 200 μg/mL bovine serum albumin (BSA), 2 mM MgCl_2_, and 1 mM DTT, along with 76 fmol (total DNA) of the topoisomer pool, and 200 nM Mlh1-FLAG-Pms1 (final concentrations). Reactions were incubated for 30 minutes at 4°C and oscillated at 20 RPM. Mlh1-FLAG-Pms1 and any associated DNA molecules were immunoprecipitated using M2 anti-FLAG magnetic beads (Pierce). Briefly, 30 μL of suspended beads (25% slurry in PBS pH 7.2, containing 0.01% Tween-20 detergent and 0.02% sodium azide) were added to each reaction and incubated for 30 min at 4°C with constant oscillation at 20 RPM. The supernatant was then separated from the beads using neodymium magnets and removed. The beads were washed with 10 μL of Buffer A three times and resuspended in 20 μL of Buffer A containing 0.96 units of Proteinase K. The material was then incubated at 37°C for 30 min with constant oscillation at 20 RPM followed by incubation at 95°C for 5 minutes to fully denature and degrade Mlh1-FLAG-Pms1 and dissociate the DNA from the beads. The denatured supernatant was then analyzed by 1% (w/v) agarose gel resolved in 1x TBE (89 mM Tris base, 89 mM boric acid, 2 mM EDTA) and stained in 1x TBE containing 1x GelRed (Biotium). Methods for quantifications are described in the Results section as well as in the supplemental information.

### DNA Topoisomerase and Gyrase Enhancement Assays

Mlh1-Pms1’s ability to stimulate *E. coli* Topoisomerase I and *E. coli* DNA Gyrase were assayed by similar methods. For the Topoisomerase I enhancement assays, 76 fmol of supercoiled pUC18 was incubated with increasing concentrations of Mlh1-Pms1 (50, 100, or 200 nM) for 10 minutes at 25°C in a buffer containing 50 mM potassium acetate, 20 mM Tris-acetate pH 7.9, 10 mM magnesium acetate, and 100 μg/mL BSA in a final volume of 20 μL. After the initial incubation, 0.05 units of Topoisomerase I (NEB) were added, and the reactions were incubated for 20 minutes at 37°C. Reactions were stopped with 0.96 units of Proteinase K, 1% SDS, and 14 mM EDTA (final concentrations), then resolved by a 1% (w/v) agarose gel, electrophoresed in 1x TBE for 75 minutes. Gels were stained in 1x TBE containing 1.6 μg/mL ethidium bromide for 30 minutes. For the Gyrase enhancement assays, pre-relaxed pUC18 was incubated for 10 minutes at 25°C in a 20 μL reaction with the same concentrations of Mlh1-Pms1 above in a buffer containing 35 mM Tris-HCl pH 7.5, 24 mM KCl, 4 mM MgCl_2_, 1 mM ATP, 2 mM DTT, 1.8 mM spermidine, 6.5% glycerol, 100 μL/mL BSA. After this initial incubation, 0.0075 units of *E. coli* DNA Gyrase (TopoGEN) were added, and the reactions were incubated for 30 minutes at 37°C. The enhancement of both enzymes was quantified by comparing the band intensities of either fully relaxed DNA, in the case of Topoisomerase I, or fully supercoiled DNA, in the case of Gyrase, relative to the input DNA concentrations for the respective set of reactions.

### UV-crosslinking assays

76 fmol of supercoiled or relaxed pUC18 was incubated with increasing concentrations of Mlh1-Pms1 and irradiated with UV light (300 nm) for 60 minutes. Reactions were irradiated at room temperature in 200 μL transparent PCR tubes (VWR #76318-804), 15 cm from the light source in a Rayonet Photochemical Reactor-200 equipped with 12 lamps outputting 8 W each (Southern New England Ultraviolet Company RPR-3000A). Following irradiation, reactions were treated with 0.96 units of Proteinase K for 15 minutes at 37°C, and products were analyzed by 1% (w/v) agarose gel resolved in 1x TAE (40 mM Tris-acetate, 1 mM EDTA) and stained with 1.5 μg/mL ethidium bromide.

### T7 Endonuclease I Enhancement Assays

Similar procedures were used to test Mlh1-Pms1’s ability to stimulate a structure-specific endonuclease on both supercoiled and nicked pUC18. The nicked pUC18 was generated by incubating supercoiled plasmid with Nt.BspQI (NEB) according to the manufacturer’s instructions, followed by heat inactivation. To measure T7 endonuclease I stimulation, DNA (3.8 nM) was added to a 20 μL reaction containing 50 mM NaCl, 10 mM Tris-HCl pH 7.9, 10 mM MgCl_2_, and 1 mM DTT (final concentrations). Where indicated, 50 nM Mlh1-Pms1 was added to reactions containing supercoiled pUC18, and 100 nM Mlh1-Pms1 was added to reactions containing nicked pUC18. Reactions were incubated at room temperature for 10 minutes to allow Mlh1-Pms1 to bind to the DNA. Subsequently, 0.5 mM ATP or ATP*γ*S was added where indicated, and the reactions were incubated at 37°C for 15 minutes. Following this, 0.2 units of T7 endonuclease I (NEB) was added to reactions with supercoiled pUC18, and 0.6 units were added to reactions with nicked pUC18, as indicated. The reactions were incubated at 37°C for 45 minutes and stopped by adding 1% SDS, 14 mM EDTA, and 0.96 units of Proteinase K (final concentrations).

The reaction products were resolved on a 1% (w/v) agarose gel containing 0.66 μg/mL ethidium bromide in 1x TAE buffer.

### ATPase Assays

ATPase activity was measured using thin layer chromatography (TLC) as previously reported (17, 39). Briefly, 10 μL reactions were prepared in a buffer containing 20 mM HEPES-KOH pH 7.5, 20 mM KCl, 1% glycerol, 2 mM MgCl_2_, 2.5 mM MnSO_4_, and 40 μg/mL BSA including 3.8 nM of either supercoiled pBR322 or pBR322 relaxed with a topoisomerase as described above, 104 μM [*γ*-^32^P]-ATP (Perkin Elmer), and 400 nM of Mlh1-Pms1 (final concentration). Reactions were incubated for 45 minutes at 37°C. ATP hydrolysis products were analyzed using polyethylenimine cellulose plates resolved in a solution of 0.8 M LiCl and 1 M formic acid for 35 minutes. Plates were phosphor-imaged using a Sapphire Biomolecular Imager (Azure). To quantify the amount of ATP hydrolysis, the density of the spot corresponding to hydrolysis product was quantified relative to the total amount of signal in each lane. The proportion of [*γ*-^32^P]-ATP hydrolyzed was then converted to pmol of ATP using the total number of pmol of ATP in the reaction.

### Electrophoretic Mobility Shift Assays

Electrophoretic mobility shift assays were used to assess DNA binding to plasmid-based substrates. Supercoiled or relaxed pUC18 DNA (76 pmol) or its derivative with a dinucleotide repeat insert was incubated with increasing concentrations of Mlh1-Pms1 in 20 μL reactions in a buffer containing 20 mM HEPES-KOH pH 7.5, 6% glycerol, 2 mM MgCl_2_, 1 mM DTT, and 200 μg/mL BSA for 20 minutes at ambient room temperature, unless otherwise indicated. Mlh1-Pms1-bound DNA was separated from unbound DNA using a 1% (w/v) agarose gel resolved in 1x TAE on ice and stained with 1.5 μg/mL ethidium bromide.

### Endonuclease Assays

Endonuclease assays were performed as previously described (13, 27, 28, 40). Briefly, 76 pmol of DNA substrate was incubated with the indicated concentrations of Mlh1-Pms1 in 20 μL reactions containing 20 mM HEPES-KOH (pH 7.5), 20 mM KCl, 1% glycerol, 2.5 mM MnSO_4_, 0.5 mM ATP, and 200 μg/mL BSA. Final concentrations of PCNA and RFC were 0.5 μM and 0.1 μM, respectively. Reactions were incubated at 37°C for 60 minutes, followed by the addition of 0.96 units of Proteinase K, 1% SDS, and 14 mM EDTA (final concentrations) to stop the reaction. Endonuclease products were analyzed by either a 1% (w/v) agarose gel containing 0.66 μg/mL ethidium bromide for supercoiled substrates or a 1% (w/v) agarose gel containing 30 mM NaCl and 2 mM EDTA when a denaturing agarose gel was used to measure nicking on relaxed or linear DNA topologies. Denaturing gels were resolved in a buffer solution containing 30 mM NaOH and 2 mM EDTA at 30 V for 2.5 hours, followed by neutralization in a 1 M Tris-HCl pH 7.5 buffer and staining with 0.5 μg/mL ethidium bromide.

To quantify the amount of supercoiled DNA nicked by Mlh1-Pms1 using native agarose gel analysis, band intensities corresponding to the supercoiled starting material and the nicked product were measured. The proportion of nicked product was calculated relative to the sum of these bands, subtracting the DNA signal at the relaxed or nicked position in the negative control lanes. To quantify the amount of relaxed or linear DNA nicked by Mlh1-Pms1 using denaturing agarose gel analysis, band intensities of the starting material were measured, and the signal loss relative to the starting material in negative controls was calculated.

### Gel Imaging

Gels were imaged on a Sapphire Biomolecular Imager (Azure) quantified using ImageJ software (NIH). Quantifications for individual assays are described under each assay. Where gel images are present, representative images are shown. Quantifications below gels refer to the selected image, averages between replicates, and standard deviations between replicates.

## RESULTS

### ATP modulates Mlh1-Pms1 activities on supercoiled DNA distinctly from relaxed DNA

Previous studies have suggested that the Mlh1-Pms1 endonuclease binds cooperatively to DNA, with multiple copies of the complex likely present at a stoichiometric excess on the DNA during mismatch repair (24–29, 36). This cooperative binding is thought to promote DNA-DNA associations, a critical activity for activating the endonuclease (27, 28), and may also induce conformational changes in DNA, as previously suggested (23, 26).

To investigate how Mlh1-Pms1 promotes and uses DNA-DNA associations, we first examined the role of DNA conformation on the protein’s affinity for DNA. Previous work shows that larger DNA molecules generally serve as better substrates for binding and activity than smaller ones (23, 27, 28), likely due to their increased flexibility and dynamic behavior in solution. If Mlh1-Pms1 associates two regions of DNA, larger substrates may provide a more favorable environment for these interactions. Moreover, previous research has demonstrated that endonuclease activity is moderately higher on supercoiled plasmid DNA compared to relaxed or linear plasmid forms (Witte *et al*, 2023). The supercoiled conformation brings helical regions closer together, potentially enhancing the DNA-DNA interactions.

To directly measure Mlh1-Pms1’s affinity for DNA as a function of supercoiling, we employed a modified high-throughput assay (41) to evaluate its binding to topoisomers of the same plasmid differing only in their writhe number, or degree of supercoiling (Figure 1A). We prepared a pool of DNA substrates by treating a pUC19 plasmid, where over 90% of the molecules were in a negatively supercoiled state, with a small amount of *E. coli* Topoisomerase I (a type I topoisomerase) for 60 seconds at 25°C followed by heat inactivation. Using this method, we generated seven distinct topoisomers, which were resolved by agarose gel electrophoresis (Figure 1B, lane 2). These topoisomers were assigned numerical identities from zero (most relaxed) to six (most supercoiled).

**Figure 1:**
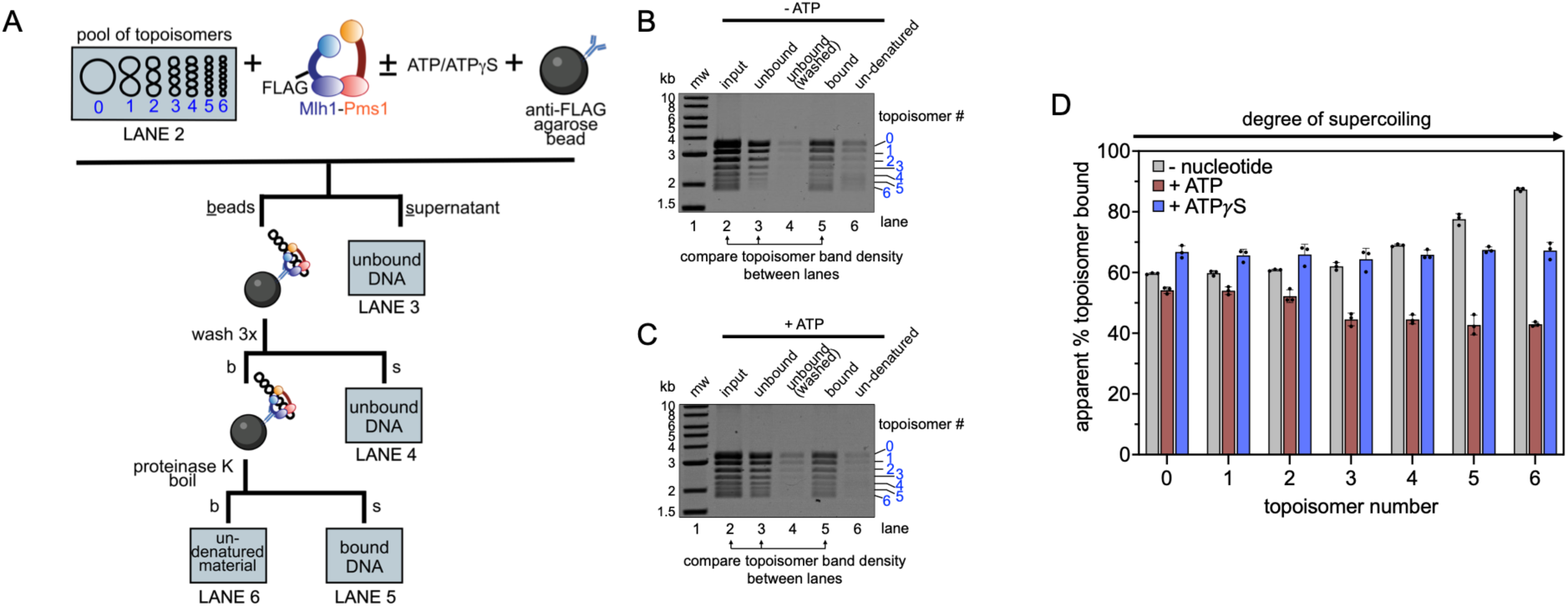
High-throughput assay measuring supercoiling density and nucleotide effects on Mlh1-Pms1 binding. (A) Modified topology-dependent binding assay reported by Litwin, *et al*. Creation of topoisomer pool and assay conditions are described in the Materials and Methods. We were able to resolve seven distinct topoisomers and assigned them arbitrary numbers 0-6 from low to high supercoiling based on relative migration in an agarose gel. The input, supernatants from each separation step, and material where denaturation was incomplete were analyzed by agarose gel. Lane numbers corresponding to analysis in panels B and C are given. (B-C) Representative agarose gel analysis of reaction material. Where included, the concentration of ATP or ATPγS was 0.5 mM. 200 nM of Mlh1-Pms1 was added in all reactions. (D) The amount of each topoisomer was calculated in the bound and unbound fractions and compared to the input. See Figure S1 for quantification details. Because a portion of the bound population resisted Proteinase K treatment and heat denaturation, the apparent amount of bound DNA expressed in the plots is input minus the sum of the unbound population for each topoisomer.

We incubated this pool of topoisomers with a FLAG-tagged variant of Mlh1-Pms1 (Mlh1-FLAG-Pms1), under conditions that do not promote endonuclease activity (i.e.—no PCNA or manganese were added). Previous studies have shown that this tag does not disrupt Mlh1-Pms1 function (16). After allowing Mlh1-FLAG-Pms1 to bind to the DNA pool, we used anti-FLAG M2 agarose beads to pull down Mlh1-FLAG-Pms1 and any associated DNA. Following several wash steps, we treated the bead-bound material with Proteinase K and then boiled it to release the Mlh1-FLAG-Pms1-bound DNA (Figure 1A). We ran all material in an agarose gel and quantified the total amount of DNA in each topoisomer band in the unbound and bound fractions after the washing steps relative to the input material to quantify the amount of each topoisomer bound by Mlh1-FLAG-Pms1 (Figure 1B-C, Figure S1 for details on quantifications).

Using this method, we found that in the absence of nucleotide, the apparent fraction of DNA bound increased with higher degrees of supercoiling (Figure 1D, grey bars). When ATP was included in the reaction, there was minimal change in the proportion of relaxed DNA bound by Mlh1-Pms1. However, the apparent proportion of supercoiled DNA bound by Mlh1-Pms1 decreased significantly (Figure 1D, compare grey bars to maroon bars for each topoisomer). This trend was observed as a function of the degree of supercoiling. When ATP*γ*S was included, Mlh1-Pms1 bound all DNA topologies to a similar extent, comparable to the binding of relaxed plasmid DNA in the absence of nucleotide (Figure 1D, blue bars). Our data suggest that the protein’s interactions with relaxed DNA are less affected by ATP, indicating that when bound to relaxed DNA, Mlh1-Pms1 may not be in a conformation to bind or hydrolyze ATP efficiently. In contrast, DNA supercoiling creates a sensitivity to ATP that either causes Mlh1-Pms1 to dissociate from the DNA or alters its conformation such that it cannot be effectively pulled down in this assay.

### Mlh1 and Pms1 ATPase Activities Play Distinct Roles in Modulating DNA Shape

To better understand the relationship between Mlh1-Pms1’s ATPase activity and its interactions on relaxed and supercoiled DNA substrates, we leveraged our previous observation that Mlh1-Pms1 can stimulate *E. coli* Topoisomerase I, a type I topoisomerase (28). Previously, we hypothesized that this stimulation was due to Mlh1-Pms1 promoting DNA-DNA associations, thereby generating a substrate more favorable for the topoisomerase. Expanding on this, we aimed to determine whether Mlh1-Pms1 could similarly influence the activity of other enzymes that alter DNA topology and to what extent ATPase activity was required for these effects.

Consistent with its stimulatory effect on *E. coli* Topoisomerase I, Mlh1-Pms1 was also able to enhance the activity of *E. coli* Gyrase, a type II topoisomerase that converts relaxed plasmids to supercoiled topologies (Figure 2A-B). Since Gyrase uses ATP hydrolysis to convert relaxed DNA into supercoiled DNA, we next tested whether ATP hydrolysis by Mlh1-Pms1 was necessary for this stimulation. For this, we used an Mlh1-Pms1 mutant (mlh1N35A-pms1N34A) that cannot bind or hydrolyze ATP in either subunit (10, 11, 17). In the presence of this mutant, we no longer observed enhancement of DNA supercoiling compared to the negative control, suggesting that Mlh1-Pms1’s ability to stimulate Gyrase activity requires ATP binding and subsequent rearrangement of the DNA (Figure 2C). When we performed this assay with mutants where ATP binding in individual subunits was inhibited (mlh1N35A-Pms1 and Mlh1-pms1N34A), we found that when ATP binding in the Mlh1 subunit was eliminated, Gyrase could still be stimulated to supercoil the DNA (Figure 2D, compare lanes 5-7 with 3). However, when ATP binding in the Pms1 subunit was abolished, Gyrase stimulation was apparently lost (Figure 2D, compare lanes 9-11 with 3), suggesting that ATPase activity in the Pms1 subunit is important for Mlh1-Pms1’s manipulation of the DNA into a conformation that stimulates Gyrase.

**Figure 2:**
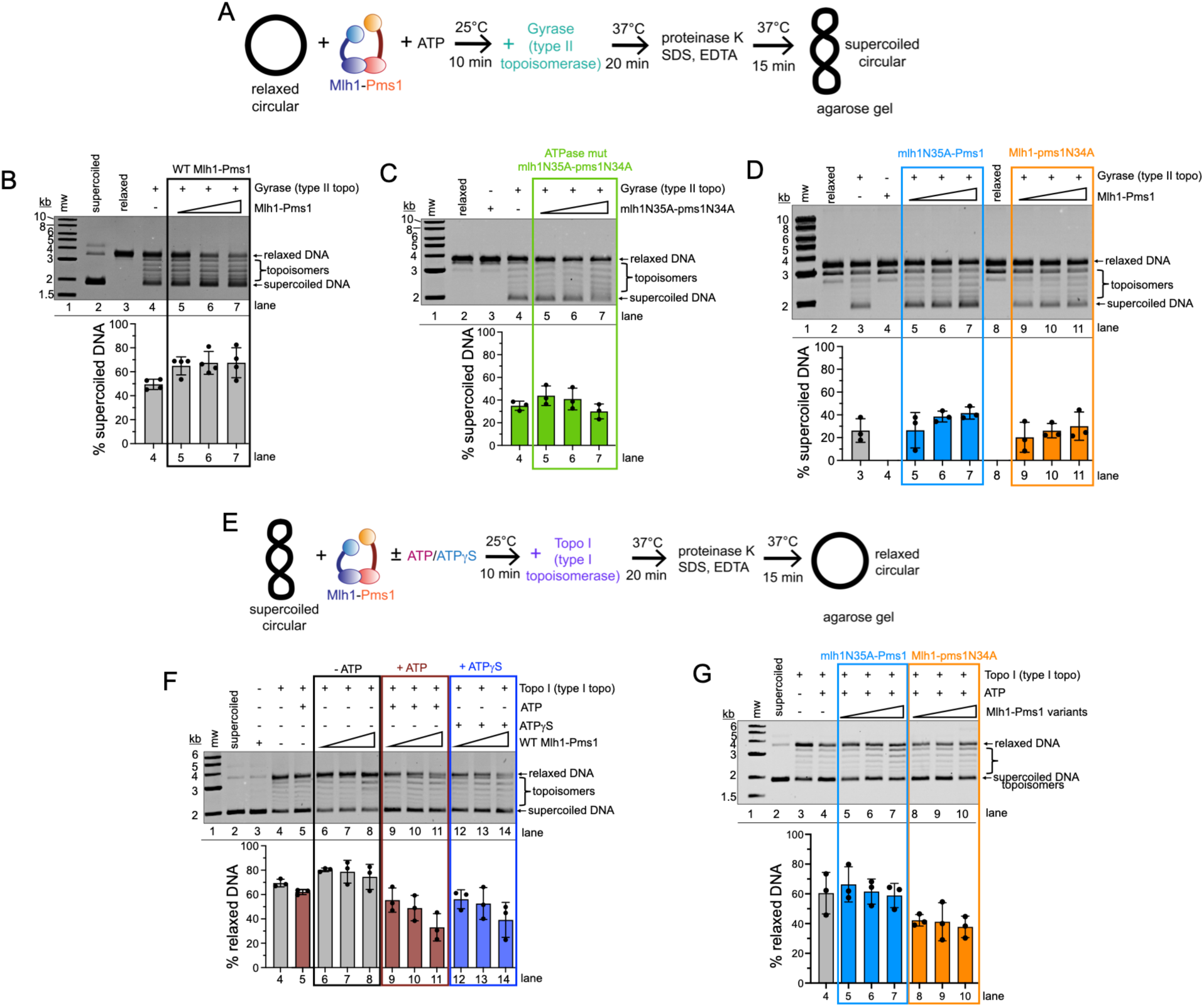
Mlh1-Pms1 enhances Topoisomerase and Gyrase, with distinct roles for each subunit. (A) Schematic for assay probing for *E. coli* Gyrase enhancement. Relaxed substrate was prepared by incubating supercoiled DNA with a topoisomerase, then inactivating the topoisomerase, see Materials and Methods for details. The relaxed circular DNA was then incubated with Mlh1-Pms1 or Mlh1-Pms1 ATPase mutants (titrated at 50, 100, or 200 nM) in the presence of 1 mM ATP, followed by the addition of *E. coli* Gyrase. Reaction products were analyzed by agarose gel. (B-D) The average proportion of supercoiled product relative to the total amount of DNA in each lane is reported for triplicate experiments. Error bars represent the standard deviation between experiments. (E) Schematic for assay probing *E. coli* Topoisomerase I enhancement. Supercoiled DNA was incubated with Mlh1-Pms1 (titrated at 50, 100, or 200 nM where indicated) in the presence or absence of 0.5 mM ATP or ATPγS, followed by the addition of Topoisomerase I (Topo I) from *E. coli*. Reaction products were analyzed by agarose gel. (F) Summary data from agarose gel analysis of assay described in panel E using wild-type Mlh1-Pms1. (G) Summary of agarose gel analysis of assay described in panel E using mlh1N35A-Pms1 or Mlh1-pms1N34A ATPase mutants. For panels F-G, the average proportion of relaxed circular DNA product relative to the total amount of DNA in each lane is reported for triplicate experiments. Error bars represent the standard deviation between experiments.

To also determine the roles of individual subunits and the effects of ATP on Mlh1-Pms1’s ability to stimulate *E. coli* Topoisomerase I, we performed the previously reported topoisomerase assay with the modification that we used 50-fold less topoisomerase to keep the degree of stimulation within the linear range, amplifying subtle effects (Figure 2E-F) (28). When either ATP or ATP*γ*S was added to this assay under these conditions, we observed a decrease in the formation of the relaxed circular DNA product compared to equivalent protein concentrations without nucleotide (Figure 2F, compare lanes 9-11 and 12-14 with 6-8). These data suggest that Mlh1-Pms1 in the ATP-bound state inhibits topoisomerase activity.

To investigate the role of each ATPase site in the enhancement of Topoisomerase I, we used the Mlh1-Pms1 mutants where ATP binding was abolished in each subunit individually. ATP was included in all reactions. In the absence of ATP binding by the Mlh1 subunit (mlh1N35A-Pms1), we observed no significant inhibition of topoisomerase or stimulation compared to the control (Figure 2G, compare lanes 5-7 with 4). Conversely, when Mlh1-Pms1 containing an ATP-binding mutation in the Pms1 subunit was used (Mlh1-pms1N34A), we observed a decrease in the proportion of relaxed DNA created by topoisomerase (Figure 2G, compare lanes 8-10 with 4), similar to that seen for the wild-type protein in the presence of ATP*γ*S.

Based on these results, we hypothesize that, on compact supercoiled DNA, ATP binding to Mlh1-Pms1—driven by the Mlh1 subunit—induces a conformational change in the protein, which in turn may alter the arrangement of the DNA bound by Mlh1-Pms1. This rearrangement may contort DNA to a conformation that is protected or inhibitory for Topoisomerase I activity. Together, our data suggest that Mlh1-Pms1 may utilize ATP binding by each subunit for distinct purposes to modulate DNA structure as measured by differences in stimulation to enzymes that modulate DNA topology.

### Mlh1-Pms1 reshapes DNA using ATP binding

To investigate how Mlh1-Pms1 alters DNA topology to stimulate Topoisomerase I and Gyrase, and how ATP influences these rearrangements, we conducted a UV-stimulated DNA-DNA crosslinking assay. In this assay, purified Mlh1-Pms1 was combined with relaxed circular plasmid DNA, and reactions were performed with or without ATP or the non-hydrolyzable analog ATP*γ*S. UV exposure at 300 nm induces inter-and intra-strand crosslinks, creating a covalent attachment between DNA regions brought into close proximity transiently by Mlh1-Pms1 binding, allowing us to take a snapshot of potential DNA reshaping, such as compaction or loop formation (Figure 3A). Using relaxed circular plasmid DNA in this assay, we observed that addition of Mlh1-Pms1 resulted in a shift to a more heterogeneous DNA population, likely reflecting Mlh1-Pms1 binding or crosslinking (Figure 3B, compare lane 4 with lanes 2 and 3). This shift was not the result of Mlh1-Pms1 nicking the DNA, because this assay was performed under conditions where Mlh1-Pms1’s endonuclease activity is not observed due to the absence of RFC or PCNA (Figure S2). After Proteinase K treatment to degrade the protein component, a new faster-migrating DNA species appeared (Figure 3B, lane 5). A small amount of this species was observed without Mlh1-Pms1 (Figure 3B, lane 3), possibly due to the plasmid’s natural conformational flexibility, but its abundance increased markedly with Mlh1-Pms1 and Proteinase K. These findings suggest Mlh1-Pms1 may promote intra-molecular DNA-DNA associations, generating a looped or more compact DNA structure.

**Figure 3.**
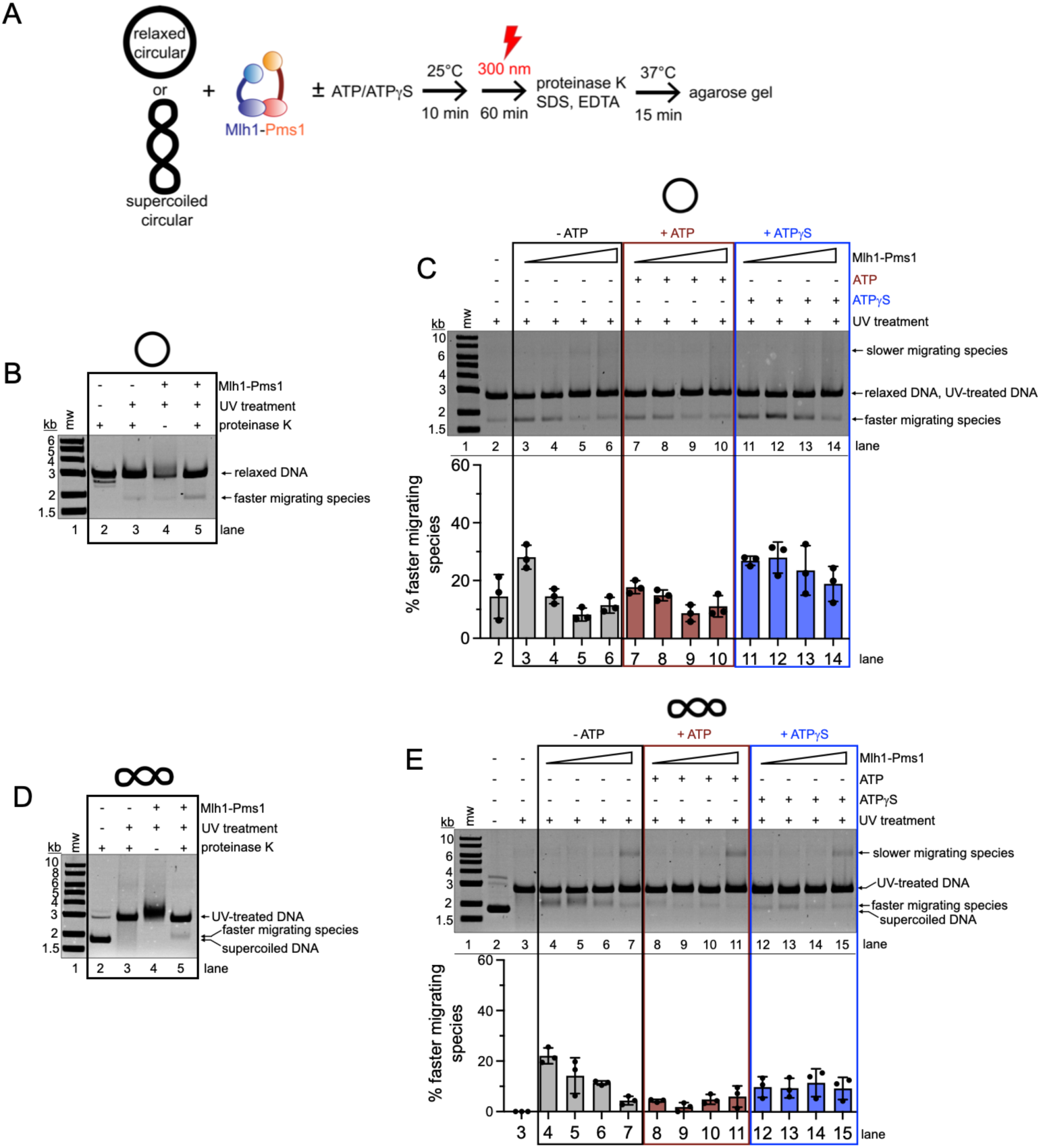
Mlh1-Pms1 rearranges DNA in an ATP-dependent mechanism. (A) Schematic for UV crosslinking assay for detecting Mlh1-Pms1-dependent DNA-DNA crosslinks. Either relaxed or supercoiled plasmid was incubated with Mlh1-Pms1 in the presence or absence of ATP or ATPγS. Reactions were then irradiated, deproteinated, and analyzed by agarose gel. See Materials and Methods for additional details. (B) Control experiment showing the effect of UV treatment on relaxed DNA with and without Mlh1-Pms1 (100 nM) and Proteinase K. (C) Mlh1-Pms1 titration and nucleotide effects on DNA-DNA crosslinking on 3.8 nM relaxed DNA. Where indicated, ATP or ATPγS was 0.5 mM. The amount of the faster migrating species was quantified relative to the total amount of DNA in each lane. The average of three replicates is reported. Error bars represent the standard deviation between experiments. (D) Control experiment showing the effect of UV treatment on supercoiled DNA with and without Mlh1-Pms1 (100 nM) and Proteinase K. (E) Identical to panel C but performed with 3.8 nM supercoiled DNA. The average of three replicates is reported. Error bars represent the standard deviation between experiments.

We hypothesized that ATP-induced conformational changes enable Mlh1-Pms1 to alter DNA shape. To test this, we repeated the assay with increasing Mlh1-Pms1 concentrations and varying nucleotide conditions. At lower Mlh1-Pms1 concentrations in the absence of nucleotide, the faster-migrating species was prominent (Figure 3C, lanes 3-4). At higher concentrations, this species diminished, and a new slower-migrating species appeared (Figure 3C, lanes 5-6), likely representing inter-molecular crosslinked plasmids. These conditions were previously seen to promote association of distinct DNA plasmids (28). The addition of ATP did not significantly change the amount of faster-migrating species compared to reactions without nucleotide, while ATP*γ*S increased its intensity (Figure 3C, lanes 11-14).

Using supercoiled plasmid DNA in this assay, we observed that UV exposure alone altered DNA migration, likely due to strand breaks (Figure 3D) (42, 43). Similar to relaxed plasmid assays, adding Mlh1-Pms1 to the reaction created a more heterogeneous population, and Proteinase K treatment revealed a faster-migrating DNA species, consistent with compacted DNA (Figure 3D, lane 5). At low Mlh1-Pms1 concentrations, this band was more prominent, while higher concentrations favored the slower-migrating species also observed with the relaxed plasmid (Figure 3E, lanes 4-5 versus 6-7). ATP reduced the faster-migrating species (Figure 3E, lanes 8-11 versus 4-7), while the slower-migrating species remained relatively unchanged, with similar results observed for ATP*γ*S (Figure 3E, lanes 12-15).

Together, these results suggest that Mlh1-Pms1 may use ATP differently on relaxed versus supercoiled DNA. We hypothesize that on relaxed DNA, ATP may be used to alter the shape of the DNA, leading to compaction. On supercoiled DNA, where the substrate is pre-compacted, ATP may drive distinct activities, such as a conformational rearrangement of the DNA and dissociation.

Based on the appearance of a faster-migrating DNA species in our crosslinking assay (Figure 3), we hypothesize that Mlh1-Pms1 compacts local DNA regions, potentially forming non-canonical structures deviating from standard B-form DNA. To better understand the nature of DNA structures potentially formed by Mlh1-Pms1, we performed an assay to assess whether Mlh1-Pms1 enhances the activity of the structure-selective T7 endonuclease I (Figure 4A). This enzyme recognizes and cleaves branched DNA structures, such as cruciform-like or looped configurations, providing insight into the topology of DNA rearranged by Mlh1-Pms1 (44–52).

**Figure 4.**
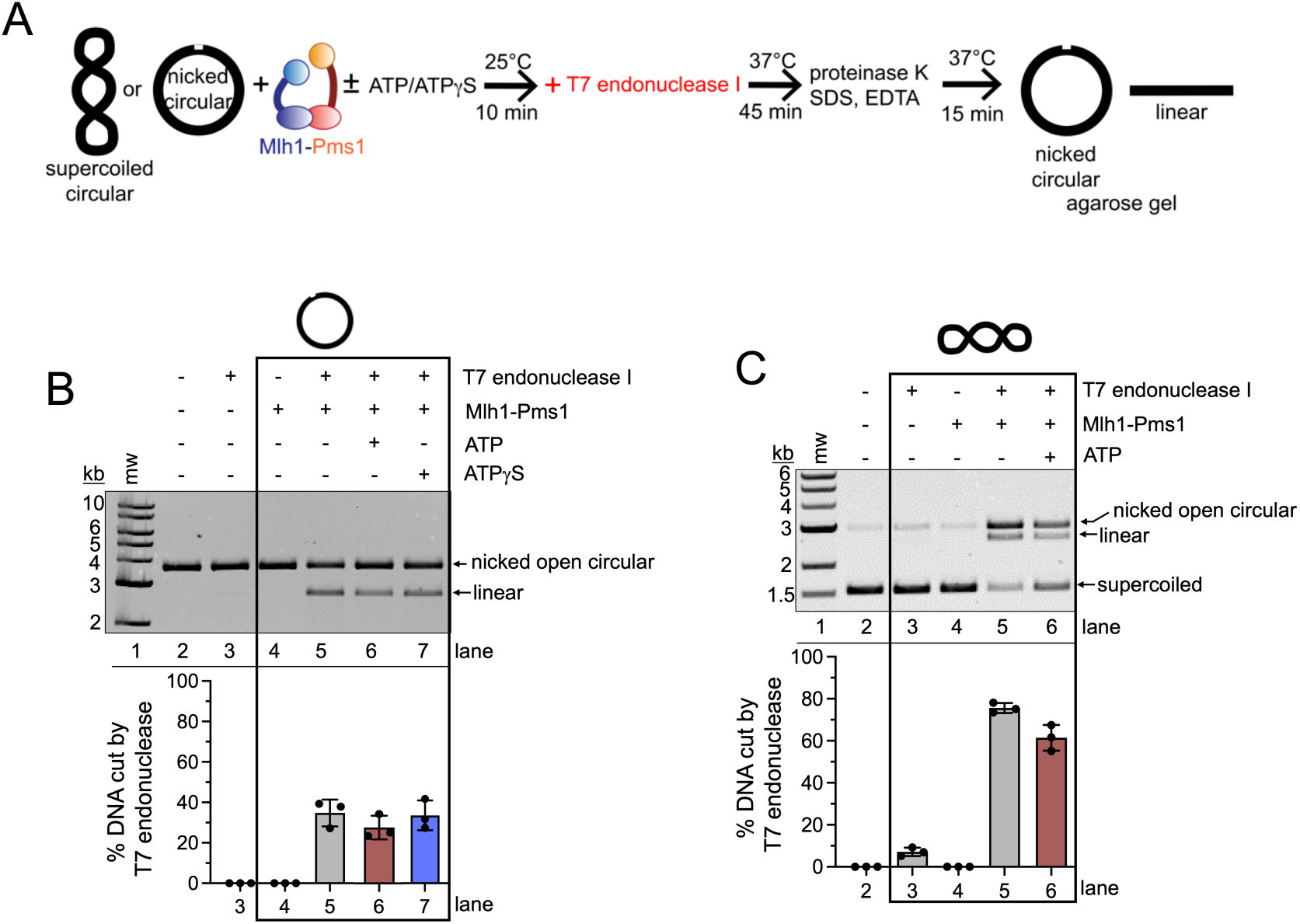
Mlh1-Pms1 rearranges DNA into a structure recognizable by a structure-selective endonuclease. (A) Schematic for structure-selective T7 endonuclease I stimulation assay in panel B and C. See Materials and Methods for additional details. (B) Agarose gel analysis of T7 endonuclease I reaction products using nicked plasmid DNA substrate generated with Nt.BspQI. Where indicated, Mlh1-Pms1 was 100 nM, ATP or ATPγS was 0.5 mM, and 0.6 units of T7 endonuclease I was added. The average amount nicked DNA converted to linear product was calculated and averaged for three replicates for each condition. This and the standard deviation between experiments is reported for each lane. (C) Agarose gel analysis of T7 endonuclease I reaction products using supercoiled DNA substrate. Where indicated, Mlh1-Pms1 was 50 nM, ATP was 0.5 mM, and 0.2 units of T7 endonuclease I was added. The average amount of supercoiled DNA converted to either nicked open circular DNA or linear product was calculated and averaged for three replicates for each condition. This and the standard deviation between experiments is reported for each lane.

We first performed the T7 endonuclease stimulation assay on a plasmid relaxed by introducing a single nick into one of the DNA strands. This approach ensured a homogenous starting material and created a substrate for T7 endonuclease I to recognize and cleave, so that we might observe stimulation of this activity. T7 endonuclease I has been shown previously to nick DNA containing a pre-existing nick, introducing a second nick and creating a double-strand break if it recognizes its substrate (53). Although our assay could not detect additional single-strand nicks that might occur from T7 endonuclease activity, it could measure double-strand breaks by observing the conversion of the open circular form to a linear form. Using this DNA substrate, we found that Mlh1-Pms1 stimulated the structure-selective activity of T7 endonuclease I, resulting in an increased linear DNA product (Figure 4B, compare lanes 3 and 5). Including ATP or ATP*γ*S had minimal effects on the stimulation (Figure 4B, lanes 6 and 7). These results suggest that Mlh1-Pms1 may compact relaxed DNA into structures resembling T7 endonuclease I substrates, but that ATP does not likely play a role in this activity.

We next incubated supercoiled DNA with Mlh1-Pms1, followed by the addition of T7 endonuclease I to assess DNA cleavage. Without Mlh1-Pms1, T7 endonuclease minimally cut the DNA (Figure 4C, compare lanes 2-3), consistent with the occasional formation of looped or cruciform-like structures due to supercoiling. Under these conditions, it is clear that Mlh1-Pms1 itself is not cleaving DNA because Mn²⁺ and RFC/PCNA, which are required for its endonuclease activity, were omitted (Figure 4B, lane 4) (13, 30, 54, 55).

When Mlh1-Pms1 was pre-bound to the DNA before adding T7 endonuclease I, a significant increase in DNA cleavage was observed, suggesting that Mlh1-Pms1 may rearrange DNA into structures that are substrates for T7 endonuclease I (Figure 4C, lane 5). When ATP was included, T7 endonuclease I cleavage was minimally reduced (Figure 4C, compare lane 6 to lane 5). This result suggests that on DNA that is supercoiled, Mlh1-Pms1 may be able to reconfigure DNA structure into a shape recognizable by the T7 endonuclease I in the absence of ATP.

### Supercoiled DNA stimulates Mlh1-Pms1’s ATPase activity more than relaxed DNA

Our data suggest that Mlh1-Pms1 may compact or reconfigure segments of DNA, bringing distant regions into proximity and potentially forming cruciform or loop-like structures. To explore whether the conformation of DNA influences Mlh1-Pms1’s ATPase activity, we performed an ATPase assay using supercoiled and relaxed plasmid substrates. Previous studies have shown that Mlh1-Pms1’s ATPase activity is stimulated by DNA (10, 13, 39), though these experiments used oligonucleotide duplexes. To specifically assess the impact of DNA topology on ATPase activity, we measured ATP hydrolysis using plasmid-based substrates and compared the results to a control reaction containing Mlh1-Pms1 alone in the absence of DNA.

We conducted two variations of this experiment. In the first, ATPase activity was measured when Mlh1-Pms1 was incubated with DNA and ATP simultaneously (Figure 5A). In the second, Mlh1-Pms1 was pre-incubated with DNA substrates for 15 minutes before ATP was added, and ATPase activity was measured (Figure 5B). Larger DNA substrates (4.3 kb pBR322) were used for these experiments, as Mlh1-Pms1 and related ATPases exhibit higher ATPase activity with larger DNA molecules (56). As expected, some ATP hydrolysis activity was observed in the absence of DNA, consistent with previous studies (Figure 5A-B, lane 1) (10, 13, 39). When Mlh1-Pms1, DNA, and ATP were added synchronously, Mlh1-Pms1’s ATPase activity was stimulated to a similar extent by both supercoiled and relaxed plasmids (Figure 5A). However, when Mlh1-Pms1 was pre-incubated with DNA before ATP addition, supercoiled DNA stimulated ATPase activity to a greater extent than relaxed DNA (Figure 5B). These results are consistent with our observations from the topoisomer pull-down assay (Figure 1) and the UV crosslinking assay (Figure 3), where ATP had a larger effect on Mlh1-Pms1’s behavior on supercoiled plasmid compared to relaxed DNA.

**Figure 5:**
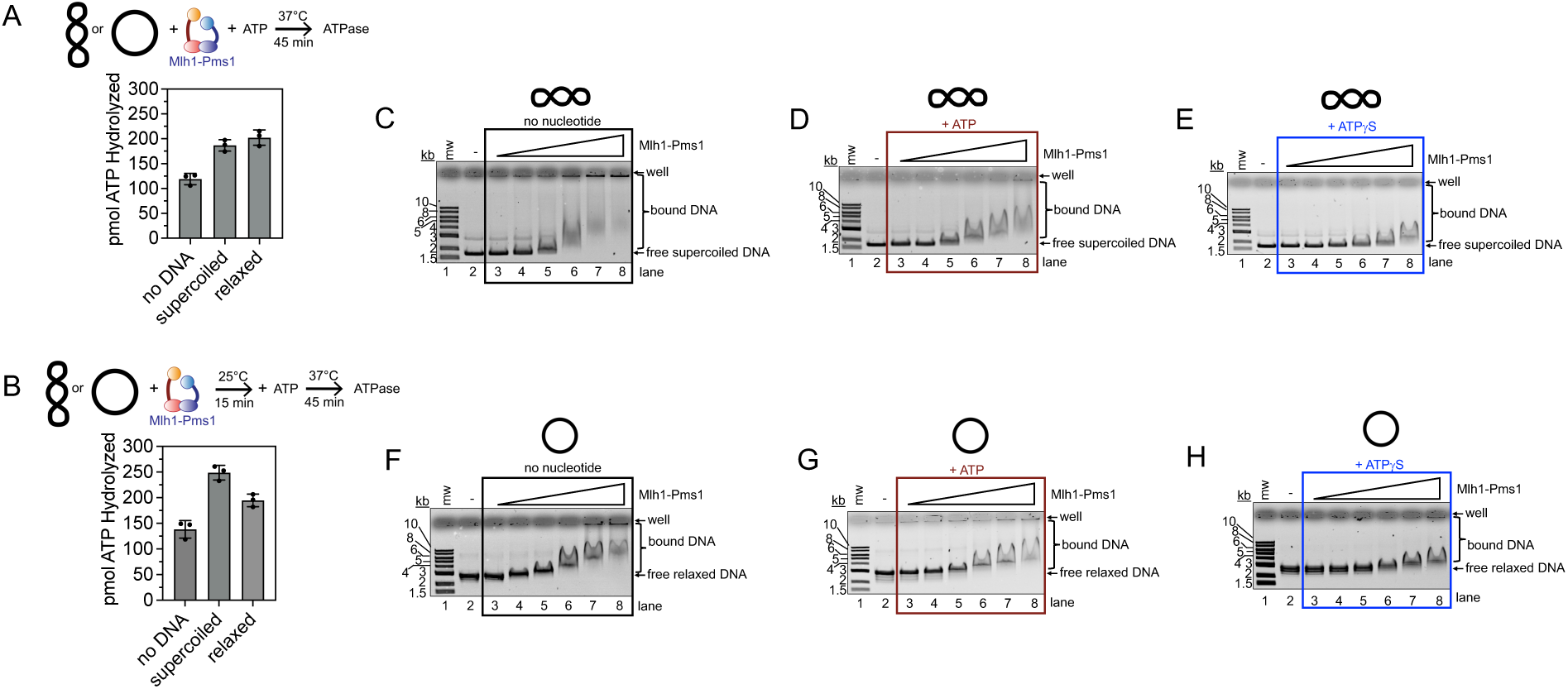
Binding to DNA in a pre-compacted state stimulates Mlh1-Pms1 ATPase. (A) ATP hydrolysis assay measuring pmol of ATP hydrolyzed in a 45 min time period. Reactions contained 400 nM Mlh1-Pms1 and 3.8 nM 4.3 kb pBR322 plasmid either supercoiled or relaxed was included where indicated. The mean and standard deviation for three replicates is reported. (B) ATP hydrolysis assays conducted by pre-binding 400 nM Mlh1-Pms1 to 3.8 nM supercoiled or relaxed pBR322 for 15 minutes, followed by the addition of ATP. Reactions were incubated at 37°C for 45 minutes. The mean and standard deviation of three replicates is reported. (C-H) Electrophoretic mobility shift assays using either supercoiled or relaxed DNA substrates at 25 mM NaCl. In lanes 3-8 for each Mlh1-Pms1 was included at 25, 50, 100, 200, 300, or 400 nM. When present ATP or ATPγS was 0.5 mM. Binding reactions were incubated at room temperature for 20 min. Experiments were conducted in triplicate, and representative gels are presented for each condition, illustrating consistent observations across all replicates.

Previous studies have suggested an interplay between Mlh1-Pms1’s ATPase activity and DNA binding, with DNA stimulating ATPase activity and ATP increasing dissociation constants for small oligonucleotide substrates (10, 13, 16). Since we observed differing levels of ATPase activity stimulation depending on the topology of the same DNA plasmid, we investigated whether Mlh1-Pms1’s affinity for different DNA topologies is affected by ATP. Using an electrophoretic mobility shift assay, we measured Mlh1-Pms1’s apparent DNA-binding affinity in the presence of ATP or ATP*γ*S compared to conditions without nucleotide.

In the absence of nucleotide, increasing concentrations of Mlh1-Pms1 caused both supercoiled and relaxed plasmids to shift to slower migrating species, consistent with protein binding (Figure 5C, F). At high Mlh1-Pms1 concentrations, a significant portion of the material shifted into the wells of the gel, likely due to Mlh1-Pms1 forming multimers on the DNA. Notably, more material shifted into the wells for supercoiled DNA than for relaxed plasmids, suggesting that supercoiled DNA is likely bound with higher affinity in the absence of ATP, consistent with the pull-down assay in Figure 1. The presence of ATP appeared to have a destabilizing effect on Mlh1-Pms1 binding to supercoiled DNA, whereas it had a minimal effect on binding to relaxed plasmids (Figure 5D, G). In contrast, the non-hydrolyzable ATP analog ATP*γ*S reduced binding to both DNA substrates (Figure 5E, H). These results suggest that DNA topology and nucleotide state both influence Mlh1-Pms1 binding.

### Non-B-form DNA structures can impede endonuclease activation

Our data suggest that Mlh1-Pms1 may alter DNA conformation. To explore this further, we focused on dinucleotide repeats, which are known to form alternative DNA structures that can significantly affect local DNA topology. We generated circular plasmid-based substrates containing dinucleotide repeat segments to assess Mlh1-Pms1’s ability to induce and respond to these DNA conformational changes (Figure 6A). Specifically, we introduced a segment of repeating AT units (p(AT)_21_) or GC units (p(GC)_22_) predicted to form cruciform or Z-form structures (57–61). These substrates were generated by ligating duplex oligonucleotides (Table S1) into EcoRI/BamHI-digested pUC18.

**Figure 6:**
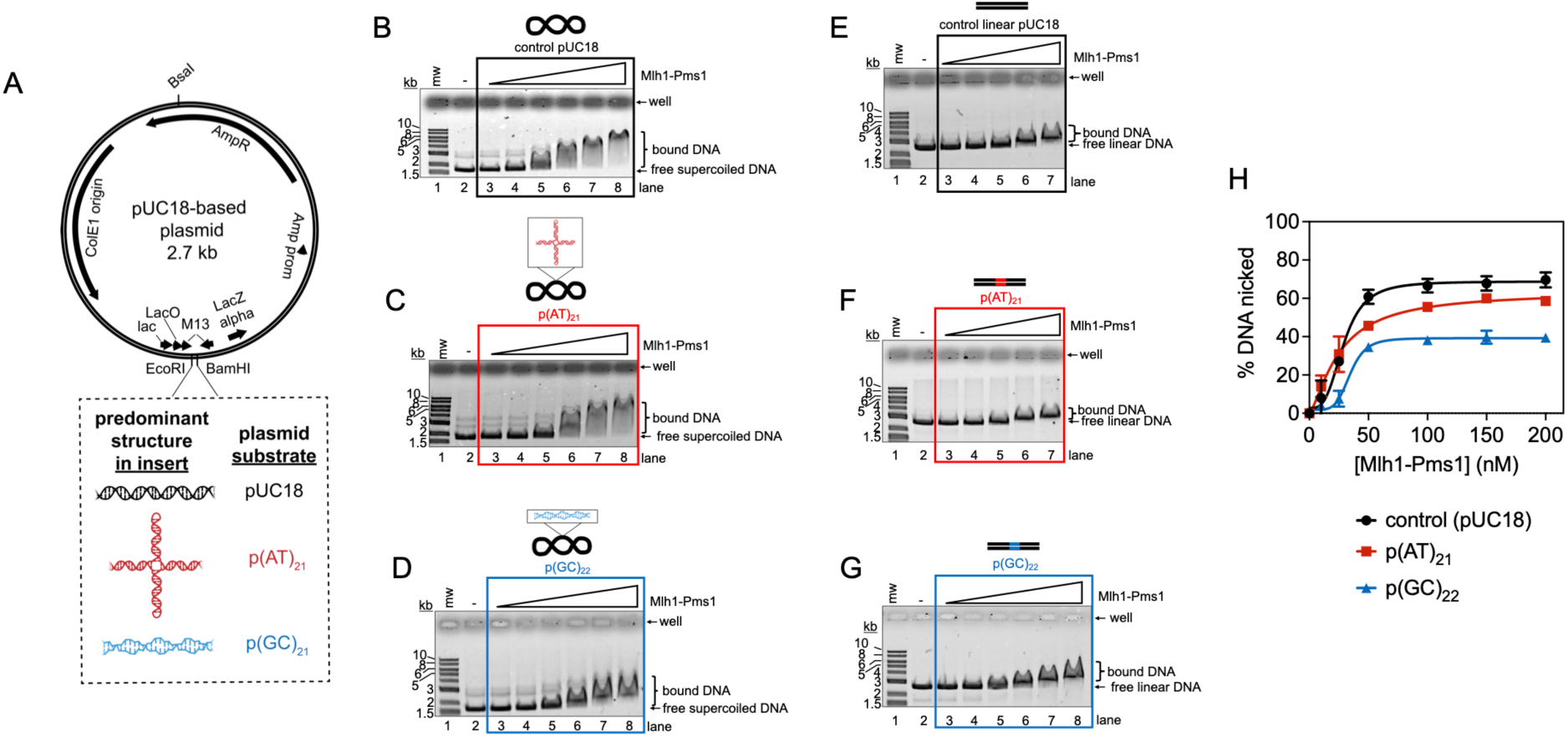
Non-B-form regions disrupt Mlh1-Pms1 activities. (A) Dinucleotide repeats were inserted into a 2.7 kb pUC18 plasmid to assess impact on Mlh1-Pms1 activities. Oligonucleotides used to construct the inserts are given in Table S1. (B-D) Electrophoretic mobility shift assays on supercoiled DNA substrates containing dinucleotide repeat regions. Where indicated, Mlh1-Pms1 concentrations are 25, 50, 100, 200, 300, 400 nM. Binding reactions were incubated at room temperature for 10 min. Experiments are performed in triplicate and representative gels are included. (E-G) DNA binding to linear substrates containing a dinucleotide repeat sequence linearized by BsaI-Hfv2, which positions the dinucleotide repeat sequence in the center of the plasmid. For Figures 6E and 6F Mlh1-Pms1 was included at 50, 100, 200, 300, 400 nM. For Figure 6G, Mlh1-Pms1 was included at 25, 50, 100, 200, 300, 400 nM. Experiments are performed in triplicate and representative gels are included. (H) Endonuclease assays on substrates containing either no repeat sequence, or a (AT)_21_ or (GC)_21_ repeat sequence. Mlh1-Pms1 was titrated at 10, 25, 50, 100, 150, 200 nM. The average proportion of supercoiled DNA converted to nicked circular product from three replicates. Error bars are the standard deviation between replicates. Data were fit to a sigmoidal function describing cooperative activity. Representative images are in Figure S4.

To verify approximately how much of each plasmid contains a cruciform structure, we used the structure-selective T7 endonuclease I, which recognizes Holliday junctions and cruciform structures. Control plasmids without dinucleotide repeat inserts were not cleaved by T7 endonuclease I in either their supercoiled or linear forms (Figure S3A, D). In contrast, the supercoiled p(AT)_21_ plasmid was efficiently cleaved by T7 endonuclease I (nearly 80% of supercoiled molecules), resulting in linearization, consistent with the formation of a cruciform structure (Figure S3B, D). When the plasmid was linearized, placing the dinucleotide repeat centrally within the linear plasmid, only ∼10% of the molecules were cleaved by T7 endonuclease I (Figure S3B, D). These findings indicate that the cruciform structure is found in a majority of molecules in the supercoiled form but is present in only a fraction of the molecules when the plasmid is linearized. For the p(GC)_22_ plasmid, ∼25% of the supercoiled molecules were cleaved by T7 endonuclease I, resulting in linearization. However, no detectable cleavage was observed in the linearized p(GC)_22_ plasmid (Figure S3C-D). This suggests that the supercoiled p(GC)_22_ plasmid may adopt a mixture of conformations, including a Z-form structure and a cruciform (60), but the cruciform structure is minimally present.

Using an electrophoretic mobility shift assay, we measured Mlh1-Pms1’s affinity for the supercoiled forms of these substrates. We found that Mlh1-Pms1 bound the p(AT)_21_ plasmid similarly to the control pUC18 plasmid (Figure 6B-C). In contrast, the p(GC)_22_ plasmid displayed a less prominent gel shift compared to the other substrates, suggesting that the p(GC)_22_ plasmid is either a lower-affinity substrate or destabilizes the Mlh1-Pms1 multimer relative to plasmids without this insert (Figure 6D). This weaker binding state appears to be dependent on the insert being in a topologically closed system, as we observed little difference in apparent affinities among the substrates when they were tested in their linear forms (Figure 6E-G). It should be noted that these binding assays were carried out for 10 minutes, as opposed to 20 minutes which was used as the incubation time for data in Figure 5 to amplify more transient effects.

A possible explanation for the weaker binding observed with the supercoiled p(GC)_22_ substrate is that this segment is predicted to form a left-handed Z-form structure. When embedded into a plasmid that is otherwise predominantly a right-handed B-form helix, this structural feature likely alters the overall biophysical parameters (e.g., twist, writhe, linking number) of the plasmid relative to the control pUC18 plasmid. Additionally, a region near the insert may become significantly deformed to maintain supercoiling in the remainder of the plasmid. This alteration in topology and conformation may result in a substrate that is less favorable for Mlh1-Pms1 binding. We also measured endonuclease activity on the supercoiled versions of these substrates.

While we observed no significant difference in the apparent affinity of Mlh1-Pms1 for the p(AT)_21_ plasmid compared to the pUC18 control plasmid, there was a reduction in the proportion of DNA nicked by Mlh1-Pms1 for the p(AT)_21_ plasmid (Figure 6H, Figure S4). The p(GC)_22_ plasmid exhibited an even greater reduction in nicking activity compared to both the pUC18 and p(AT)_21_ plasmids. Together, these data suggest that structures formed by dinucleotide repeats, such as cruciform and Z-form segments, alter local DNA topology in ways that impact Mlh1-Pms1’s affinity for and dynamics on DNA, which are critical for its function.

## DISCUSSION

Our data suggest a model in which Mlh1-Pms1 binds to relaxed DNA plasmid and leverages its ability to form oligomers and promote DNA-DNA associations to bend or rearrange the DNA into a structure where distant regions of the DNA helix are brought into close proximity (Figure 7A).

**Figure 7:**
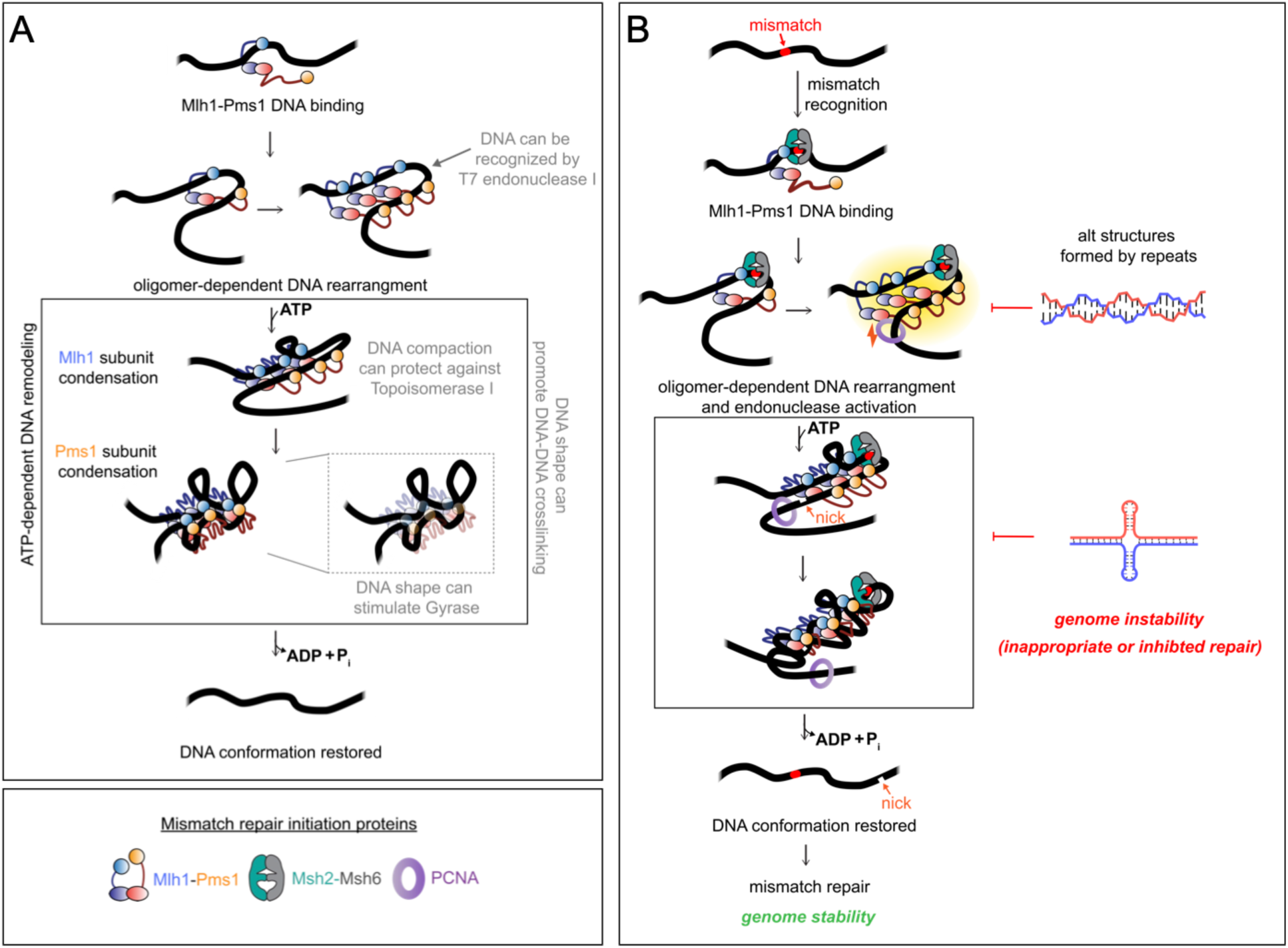
Models for Mlh1-Pms1 DNA rearrangements. (A) Model for Mlh1-Pms1 activities observed in this study. DNA-DNA associations can occur in the absence of ATP, but ATP binding by Mlh1-Pms1 affects DNA conformation as evidenced enzyme stimulation assays. The resultant DNA conformation may be highly compact, with regions resembling supercoiled (inset), which can stimulate ATP hydrolysis and Mlh1-Pms1 dissociation or sliding on the substrate. (B) After mismatch detection by Msh2-Msh6, Mlh1-Pms1 is recruited to DNA, altering its conformation via oligomerization to bring regions into proximity. Following endonuclease activity and ATP hydrolysis, the Mlh1-Pms1-induced DNA conformational change reverts. DNA structures deviating from B-form can partially inhibit key steps of this process.

This process is independent of ATP and results in a proximity sufficient to facilitate the formation of some DNA-DNA crosslinks (Figure 3C), stimulate *E. coli* Topoisomerase I (Figure 2F) (28), and create a DNA shape resembling the cruciform or loop-like substrate for the structure-selective T7 endonuclease I (Figure 4B) (44–52).

Once bound and bent, Mlh1-Pms1 can use ATP binding to change its own conformation and further rearrange the DNA substrate. We hypothesize that in the presence of ATP, condensation of the intrinsically disordered region of the Mlh1 subunit (12) facilitates DNA compaction as part of a compensatory process (Figure 7A). This is consistent with our DNA-DNA crosslinking assay, where the inclusion of ATP*γ*S increased the intensity of the faster-migrating crosslinked band on relaxed plasmid DNA (Figure 3C). This DNA compaction, initially facilitated by Mlh1, also explains our findings in Figures 2F-G, where ATP binding by Mlh1 was necessary to inhibit *E. coli* Topoisomerase I. A possible explanation for this inhibition is that DNA compaction and contortion by Mlh1-Pms1, driven by Mlh1, creates a DNA conformation in the plasmid that is inaccessible or exhibits low affinity for the topoisomerase. ATP binding by the Pms1 subunit induces further DNA rearrangement and compaction, potentially bringing regions of the DNA that were distant in the absence of Mlh1-Pms1 into even closer proximity. This is supported by our observation that ATP binding by the Pms1 subunit is necessary to stimulate *E. coli* Gyrase in our assay (Figure 2D).

These results align with previous kinetic studies using truncated amino-terminal domains of the Mlh1 and Pms1 subunits, which house the ATPase active sites. These studies demonstrated that the Mlh1 subunit has a lower K_m_ in ATPase assays compared to Pms1, while the Pms1 subunit exhibits a higher k_cat_ (17). This is also consistent with atomic force microscopy data (12), which suggest that the intrinsically disordered regions of Mlh1-Pms1—connecting the amino-terminal ATPase domains (primarily responsible for DNA binding; (6, 7)) to the globular carboxy-terminal domains—can collapse independently. This results in structures where one subunit is in a condensed form while the other is extended.

The compact DNA shape generated by ATP-bound Mlh1-Pms1 may not be supercoiled per se; however, compaction may induce loop formation, explaining why supercoiled DNA is a higher-affinity substrate (Figures 1 and 5). This compact shape also stimulates Mlh1-Pms1’s ATPase activity, which can promote protein dissociation from DNA or transition the protein into an extended conformation that slides along the DNA (8, 32, 62). This is supported by our data showing reduced Mlh1-Pms1 pull-down on supercoiled plasmids relative to other topologies in the presence of ATP (Figure 1D), increased ATPase activity when Mlh1-Pms1 is pre-bound to supercoiled DNA (Figure 5B), and apparent ATP-induced DNA dissociation observed in a gel shift assay with supercoiled plasmid (Figure 5D). Additionally, our DNA-DNA crosslinking data show that the inclusion of ATP decreases the intensity of the faster-migrating crosslinked band when supercoiled plasmids are used (Figure 3E). These findings are consistent with previous studies indicating that supercoiled plasmids better support endonuclease activity *in vitro* compared to relaxed plasmids and that ATP promotes Mlh1-Pms1 endonuclease recycling (13, 22, 28).

These findings also suggest that this DNA length may be critical for facilitating the compaction and rearrangement required to recycle Mlh1-Pms1 to observe nicking of a majority of DNA molecules in *in vitro* endonuclease assays. Supporting this hypothesis, are assays showing that perturbations altering DNA’s biophysical properties apparently inhibit endonuclease activity (Figure 6 and (28)). Additionally, this model is consistent with biochemical assays that require plasmid DNA or large linear DNA fragments exceeding the persistence length of DNA to observe reconstituted mismatch repair or Mlh1-Pms1/PMS2 endonuclease activity (27, 28).

In mismatch repair, our data suggest a model where mismatch detection by Msh2-Msh6 or Msh2-Msh3 leads to the recruitment of Mlh1-Pms1, which forms an oligomeric complex on the DNA and bends or rearranges the substrate (Figure 7B). This DNA shape may serve to restrain the Msh complex near the mismatch, as previously suggested (26, 32–35). Such restraint could protect the mismatched DNA from additional damage and prevent nucleosome deposition until repair is completed.

Previous work has shown that endonuclease activity by Mlh1-Pms1 and its homologs can occur independently of ATP (13, 21, 22). However, this activity is activated by the formation of DNA-DNA associations (27, 28), suggesting that interaction with PCNA and endonuclease activation are facilitated by the DNA conformation generated in the absence of ATP.

Following endonuclease activity, we hypothesize that the Mlh1-Pms1 complex binds ATP and undergoes a conformational rearrangement, which compacts the DNA and stimulates its ATPase activity. This ATPase activity and the associated DNA compaction may generate the free energy required to release Mlh1-Pms1 from the repair site. Interactions between Mlh1-Pms1 and PCNA further enhance ATPase activity (39), likely additionally stimulating the conformational changes necessary for dissociation or disengagement.

The DNA compaction mediated by Mlh1-Pms1 may also function as a recruitment mechanism for Exo1 or other mismatch removal factors, while simultaneously protecting the DNA from inappropriate processing. Such a mechanism would ensure the integrity of the repair site and prepare it for downstream strand removal and repair. Ultimately, the ATPase-driven release of Mlh1-Pms1 can restore the DNA conformation, facilitating efficient mismatch removal and repair.

The highly compacted DNA structure generated by Mlh1-Pms1 is also supported by atomic force microscopy data using human homologs, which reveal a large footprint of nucleotide occlusion near mismatches when MSH2-MSH6, MLH1-PMS2, and ATP are included. In these assays, “missing lengths of DNA” were observed, which could be indicative of compaction (26). These lengths ranged from approximately 50 to 300 base pairs which aligns with earlier findings showing that Mlh1-Pms1/PMS2 can nick DNA at similar distances from a mismatch (30, 31).

Our results may explain why Mlh1-Pms1 ATP-binding mutants are associated with high mutation rates *in vivo* (17, 18). Without the DNA contortions created by ATP binding, Mlh1-Pms1 may become trapped on DNA, as also suggested in exonuclease protection assays performed with ATP binding mutants (22). A trapped Mlh1-Pms1 complex could ultimately prevent mismatch removal and repair. The inability to transition to ATP-bound conformations could inhibit essential protein recycling steps, leading to defective repair and increased genomic instability.

Our findings also underscore the critical role of Mlh1-Pms1’s intrinsically disordered regions in mismatch repair (13–16). These regions mediate large conformational changes in response to ATP, and our data suggest that these changes not only alter the protein’s structure but also reshape the DNA to which Mlh1-Pms1 is bound. This process appears to be essential for releasing Mlh1-Pms1 from the DNA. Past work has demonstrated that the intrinsically disordered regions are critical for endonuclease activity *in vitro* (13–15). A plausible explanation is that for robust endonuclease activity to occur *in vitro*, Mlh1-Pms1 must nick the DNA, release the substrate, and recycle to nick additional DNA molecules in the population. Our data suggest that if these ATP-driven DNA conformational changes do not occur, Mlh1-Pms1 may fail to recycle. Defects in endonuclease activity observed in studies where the intrinsically disordered regions were altered may result from failures in this recycling or an inability to form the initial DNA structure necessary to promote DNA-DNA associations critical for repair.

Our data suggest that changes to the DNA sequence that introduce nucleic acid structures resistant to manipulation or that alter topology, such as cruciform or Z-form structures, may create substrates that are ineffectively bound or reshaped by Mlh1-Pms1 (Figure 7B). For the cruciform-forming structure, for instance, although this substrate was bound efficiently, it was less effectively nicked by Mlh1-Pms1 (Figure 6H). We hypothesize that this substrate may not be able to be manipulated in ATP-driven steps that occur after the substrate is nicked that could serve to recycle the endonuclease to nick additional substrates, explaining the apparent inhibition.

Our data point to a potential additional mechanism for genomic instability. Microsatellite regions, which consist of repeating nucleotide units, are particularly prone to adopting alternative or non-B-form structures (63, 64). These regions are hotspots for genomic instability and are commonly used as biomarkers for mismatch repair defects and associated cancers (65–68). It has been hypothesized that the repetitive nature of microsatellite sequences makes them susceptible to DNA polymerase errors and slippage, resulting in frequent mismatches and an increased demand for mismatch repair activity (66, 69). However, our data suggest that these sequences may be prone to instability not only due to an increase in errors that escape polymerase proofreading, but that mismatch repair itself may be less effective in these regions. The formation of non-B-form structures within microsatellites could interfere with Mlh1-Pms1/PMS2 activity, further contributing to genomic instability.

In addition to its evolutionary origins in DNA mismatch repair, MutL homolog complexes have been implicated in meiosis in eukaryotes, where they act on branched DNA intermediates to facilitate the formation of crossovers between chromosomes and in trinucleotide repeat instability where loop-out structures are present (70, 71). These complexes—Mlh1-Mlh2 in yeast (MLH1-PMS1 in humans and mice) and Mlh1-Mlh3—are heterodimeric and share the Mlh1 subunit. Although, our findings suggest that the mismatch repair-specific subunit Pms1 (yeast nomenclature) plays a critical role in DNA compaction and rearrangement, yeast Mlh1-Mlh3 has been shown to exhibit many biochemical features similar to those of Mlh1-Pms1, including the high degree of cooperativity and ATP-mediated conformational changes (12, 27, 28, 72). Future studies will be necessary to determine whether Mlh1-Mlh3 or Mlh1-Mlh2/PMS1 reshape DNA in a manner similar to Mlh1-Pms1/PMS2 and whether they use ATP in a comparable process.

## DATA AVAILABILITY

The data underlying this article are available in the article and in its supplementary material.

## SUPPLEMENTARY DATA

Supplementary Data are available as a separate PDF.

## AUTHOR CONTRIBUTIONS

Bryce W. Collingwood: Conceptualization, Investigation, Methodology, Writing—original draft. Amruta N. Bhalkar: Conceptualization, Investigation, Validation, Methodology, Writing—review & editing. Carol M. Manhart: Conceptualization, Methodology, Writing—original draft, Project administration, Resources, Supervision, Funding acquisition.

## Supporting information

Supplementary Data for Collingwood et al

## ACKNOWLEDGEMENTS

We thank all members of the Manhart lab, Allen Nicholson, and Eric Alani for helpful discussions about the work in this manuscript. We also thank the Department of Chemistry and the College of Science and Technology at Temple University for their institutional support. The content of this work is solely the responsibility of the authors and does not necessarily represent the official views of the National Institutes of Health or any other entity. The funders had no role in the study design, data collection and analysis, decision to publish, or preparation of the manuscript.

## FUNDING

National Institute of General Medical Sciences [R35GM142651 to C.M.M.].

## CONFLICT OF INTEREST STATEMENT

The content of this work is solely the responsibility of the authors and does not necessarily represent the official views of the National Institutes of Health. The funders had no role in the study design, data collection and analysis, decision to publish or preparation of the manuscript.

## Notes

### Competing Interest Statement

The authors have declared no competing interest.

