## Supplementary Data for Collingwood et al for "The mismatch repair factor Mlh1-Pms1 uses ATP to compact and remodel DNA"

**SUPPLEMENTARY TABLE**

| Oligonucleotide Number | Substrate | Oligonucleotide sequence (5’ to 3’) |
|---|---|---|
| CMO336 | $p(AT)_{21}$ | GATCCGCTCTTCGATCATATATATATATATATATATATAT ATATATATATATATATATGCTCTTCAGG |
| CMO337 | $p(AT)_{21}$ | AATTCCTGAAGAGCTATATATATATATATATATATATATA TATATATATATATATAGATCGAAGAGCG |
| CMO398 | $p(GC)_{22}$ | GATCCGCTCTTCGATCGCGCGCGCGCGCGCGCGCGCGCGC GCGCGCGCGCGCGCGCGCGCTCTTCAGG |
| CMO399 | $p(GC)_{22}$ | AATTCCTGAAGAGCCGCGCGCGCGCGCGCGCGCGCGCGCG CGCGCGCGCGCGCGCGGATCGAAGAGCG |

**Table S1.** Oligonucleotides used to construct plasmid substrates containing non-B-form segments. All oligonucleotides were obtained commercially from Integrated DNA Technologies. See Materials and Methods for details on substrate construction.

### SUPPLEMENTARY FIGURES

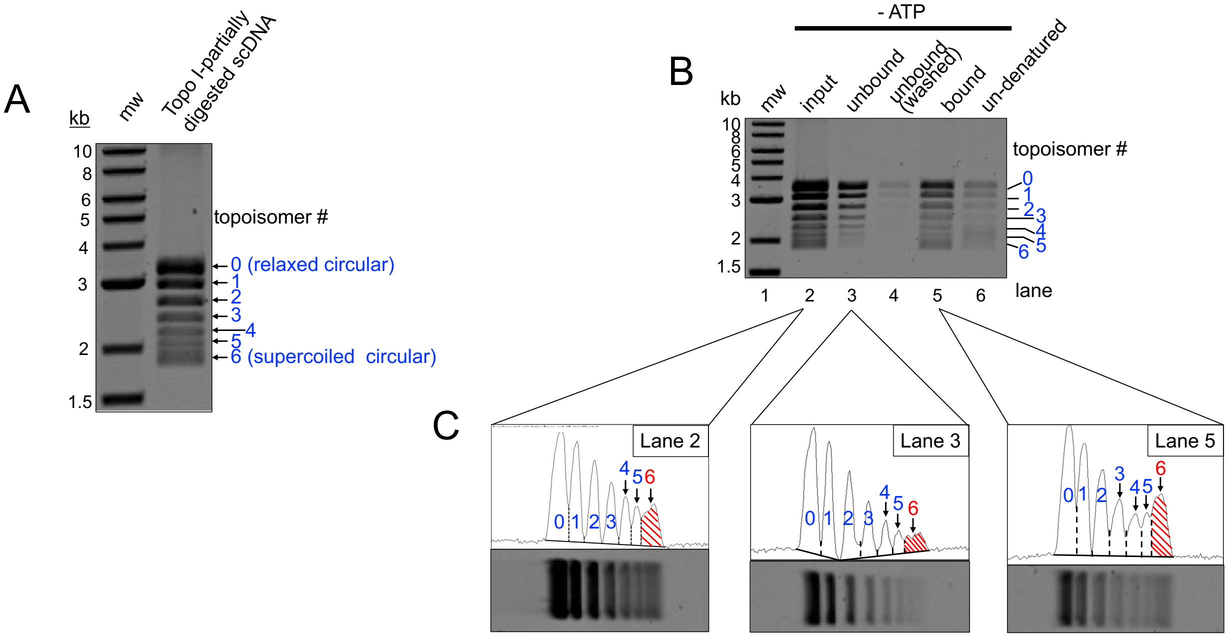


**Figure S1: Quantifications for high-throughput assay measuring supercoiling density and nucleotide effects on Mlh1-Pms1 binding.** (A) 2.7 kb pUC19 was partially digested with *E. coli* Topoisomerase I to generate a distribution of DNA topoisomers. Seven distinct bands are visible when analyzed by agarose gel. We arbitrarily numbered them 0 through 6, with zero being the most relaxed and six being the most supercoiled. This pool is the input for reactions in Figure 4. (B) Gel also shown in Figure 4B as an example of the assay and how quantifications were performed. (C) The quantification of topoisomer #6 is shown as an example (highlighted in red). Each lane was analyzed using ImageJ, where the bands corresponding to each topoisomer were identified (separated by dotted lines) and quantified. The total amount of each topoisomer in lanes 3–6 was confirmed to be equal to the amount present in the input (lane 2). For the bound population in lane 5, the density of each topoisomer band was calculated relative to the input (lane 2). Similarly, the amount of unbound topoisomer in lanes 3 and 4 (with lane 3 shown as an example) was calculated relative to the input. Because some topoisomer remained bound to the beads after denaturation and Proteinase K treatment (as seen in lane 6), the final data is expressed as the input amount minus the sum of the unbound fraction for each topoisomer. Note that the example peaks from ImageJ were resized to be visible here. The y-axes vary per lane.

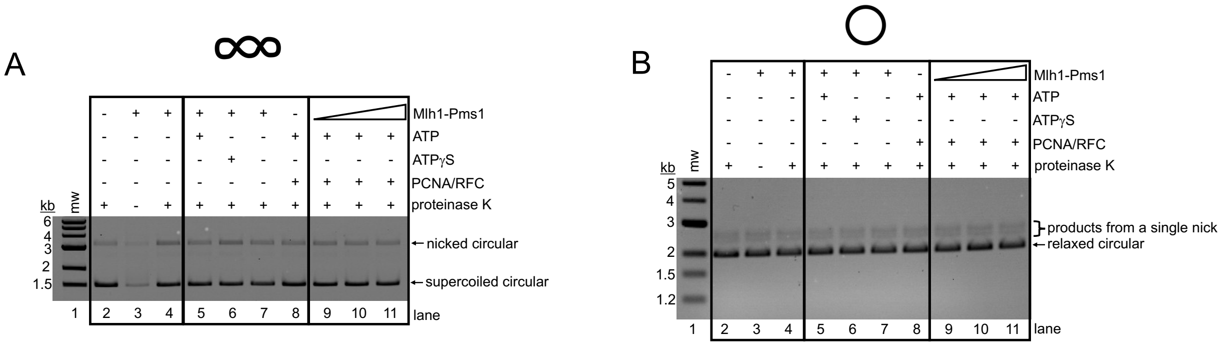


**Figure S2: Endonuclease assay confirming that conditions used for UV DNA-DNA crosslinking did not permit endonuclease activity.** (A) A supercoiled 2.7 kb pUC18 plasmid (3.8 nM) was incubated at 37ºC for 60 minutes with no UV light exposure in the same reaction conditions reported in Figure 1. Where indicated, Mlh1-Pms1 was included at 200 nM, ATP was included at 0.5 mM, PCNA was included at 0.5 mM, RFC was included at 0.1 mM, and Proteinase K was 0.96 units (final concentrations). Reaction products were analyzed by a native agarose gel and endonuclease activity is measured as conversion of supercoiled circular DNA to nicked circular. Nicked products did not increase in intensity above the negative control lanes for reactions including Mlh1-Pms1 even in the presence of RFC/PCNA and ATP due to the absence of $MnSO_4$. (B) A relaxed 2.7 kb pUC18 plasmid (3.8 nM) was tested for endonuclease activity using the same conditions as in panel A. Reaction products were analyzed by an alkaline agarose gel (described in the Materials and Methods) and endonuclease activity is measured as conversion of the relaxed circular substrate to a linear single strand fragment and a closed circular single strand fragment if a single nick occurs, or degradation of the starting material if multiple nicks occur. Nicked products did not increase in intensity above the negative control lanes for reactions including Mlh1-Pms1 even in the presence of RFC/PCNA and ATP due to the absence of $MnSO_4$ on this substrate.

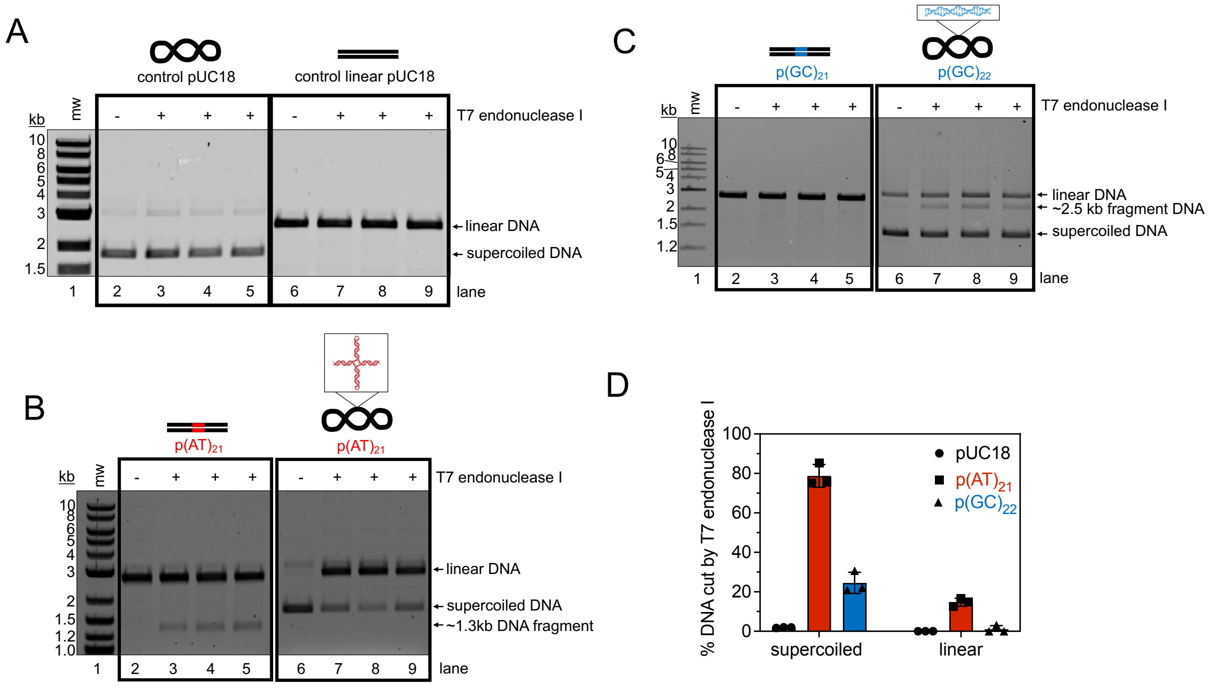


**Figure S3: Characterization of plasmid substrates containing non-B-form inserts.** (A-C) T7 endonuclease digests of plasmids containing the sequences described in Figure 6A. 3.8 nM of both supercoiled and linear forms of each DNA substrate were incubated with 1 unit of T7 endonuclease I and compared to reactions not treated with T7 endonuclease I. Where linear, the plasmids were linearized with BsaI-Hfv2. Lanes 3-5 and 6-9 of each are triplicate experiments (D) The average percent DNA cut by T7 endonuclease I and standard deviation for each plasmid in both supercoiled and linear forms compiled from (A-C).

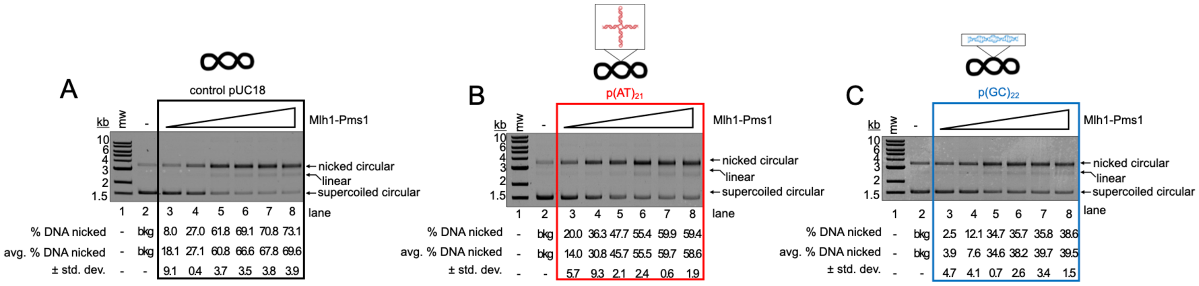


**Figure S4: Endonuclease assay on supercoiled plasmids with non-B-form inserts.** Endonuclease assays on substrates containing either no repeat sequence, or a $(AT)_{21}$ or $(GC)_{21}$ repeat sequence. Where indicated, Mlh1-Pms1 concentrations are 10, 25, 50, 100, 150, 200 nM. The average proportion of supercoiled DNA converted to nicked circular product from three replicates as well as the standard deviation between replicates are reported.
